# Design to Data for Mutant of β-Glucosidase B from *Paenibacillus polymyxa*: G23S

**DOI:** 10.64898/2026.04.27.721118

**Authors:** Amanda O’Donnell, Ghazia Abbas

## Abstract

β-glucosidase (BglB) from *Paenibacillus polymyxa* was mutated (G23S, Rosetta/Foldit numbering; G26S, conventional numbering) to assess structural and functional changes. Foldit modeling and prior Design 2 Data (D2D) database results led us to hypothesize that this mutation would increase substrate binding affinity and catalytic efficiency, with a moderate reduction in thermal stability. The mutant protein was expressed, purified, and analyzed using kinetics and thermal stability assays. Relative to the wild-type (WT), G23S exhibited a similar binding affinity (similar K_m_), an approximately 2-fold increase in turnover number (k_cat_) and catalytic efficiency (k_cat_/K_m_), an almost 14-fold increase in maximum reaction velocity (V_max_) and a slight decrease in thermostability (T_50_). The results largely support the hypothesis, indicating that changes in residue 23 can enhance catalytic power while minimally compromising stability.

## BACKGROUND

Predicting how mutations impact protein structure and function is crucial for protein engineering. Software programs like Foldit use established algorithms to predict structural and energy changes but are not entirely accurate. (1, 2) The Design to Data (D2D) project aims to use experimental data to fill in the knowledge gaps of how mutations affect proteins. (3) With a large database and a standard protocol, researchers can link mutations to biochemical parameters; the results can then improve protein software algorithms. Accurate structure to function prediction serves as the foundation of a variety of scientific and medical applications, including protein engineering, drug design, and interpretations of mutations and diseases.

β-glucosidase (BglB), part of the glycoside hydrolase (GH) family from *Paenibacillus polymyxa*, is well-characterized with substantial structural and kinetics data. The hydrolase degrades cellulose, cleaving β-glycosidic linkages; its activity is determined by its overall conformation and the chemical properties of both active site and surrounding residues. (4) The BglB structure is an (β/α)_8_ barrel with an active site in a cavity, and experimental results show that single amino acid substitutions can significantly affect its folding, conformation, and catalytic activity. (5) Thus, BglB is an ideal system for gathering structure to function data based on different point mutations.

Residue positions are reported using Rosetta/Foldit numbering, which is three amino acids shorter than the conventional numbering system. Rosetta/Foldit numbering does not include the N-Terminal fragment (NTF), so G23S in Rosetta/Foldit is G26S in conventional numbering.

BglB performs hydrolysis by a double-displacement reaction with a covalent glycosyl-enzyme intermediate. (6) Two glutamate residues (E164 and E353) and a water molecule carry out the reaction illustrated in Figure 1. Many other residues are involved in substrate binding and alignment throughout the active site and subsites, which are shaped by surrounding loops that link β/α segments. (4) Foldit modeling shown in Figure 2 predicts that the G23S mutation does not substantially alter overall protein conformation but introduces an additional hydrogen bond between S23 and E26 (blue and white line), potentially altering substrate binding and catalysis. The Foldit score also increased from -1089.697 to -1085.659, suggesting a moderate decrease in protein stability. Figure 3 shows the mutation’s location relative to other secondary structures and E26: residues 23 and 26 lie in a surface-exposed loop distal from the active site linking a β-strand to an α-helix that contains Q19. Our study investigates how the G23S substitution affects the structure, kinetics, and thermal stability of BglB.

**Figure 1.**
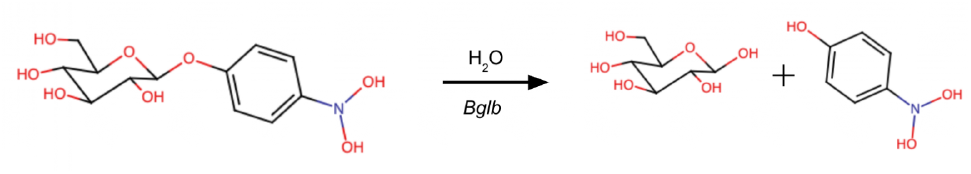
Hydrolysis reaction performed by BglB. The enzyme catalyzes the hydrolysis of β-glycosidic linkages in glycosides.

**Figure 2.**
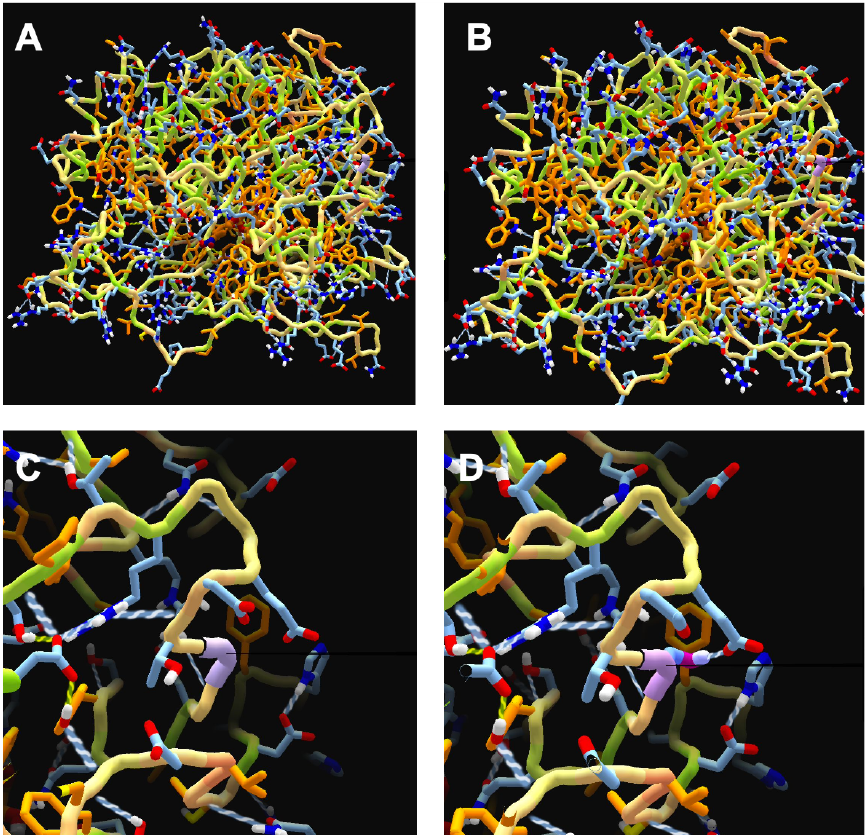
Structural comparison of wild-type (WT, panels A and C) and mutant (G23S, panels B and D) BglB modeled using Foldit. In each panel, residue 23 is highlighted in purple. Panels C and D show zoomed views of the glycine (WT) and serine (G23S) residue sites, respectively. The WT protein had an energy score of -1089.697 and a residue score of -0.859. The G23S mutant, after minimization and repack/reshake, had an energy score of -1085.659 and a residue score of 1.620. Protein and residue scores are displayed at the top and in the right-hand inset of each panel, respectively. In panel D, an additional blue and white dashed line indicates a hydrogen bond between S23 and E26 not present in the WT structure.

**Figure 3.**
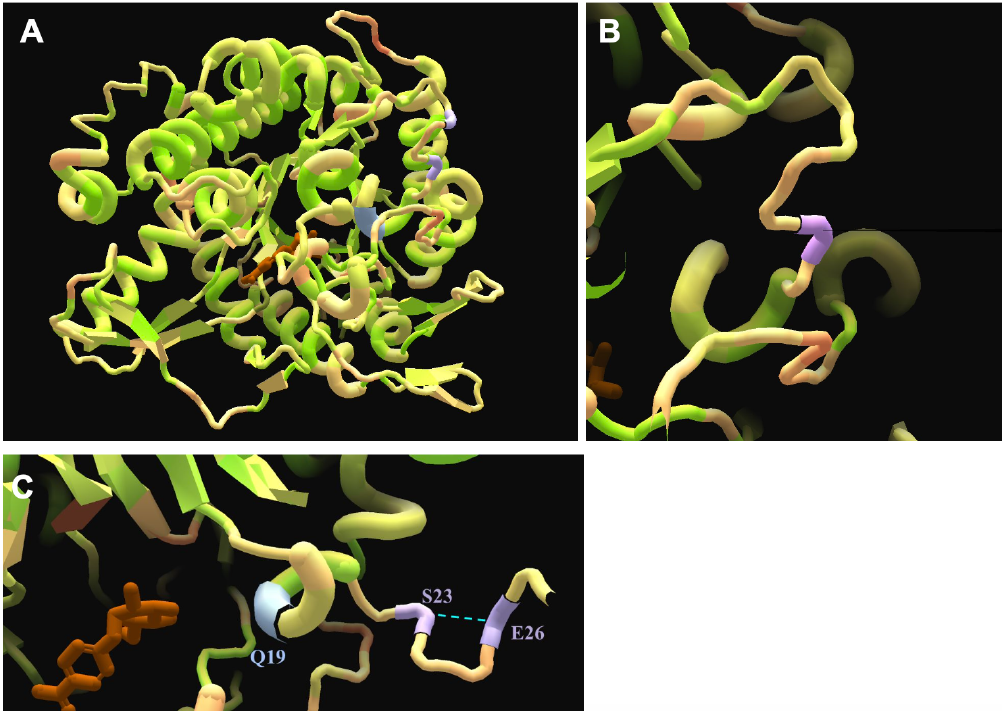
The secondary structure context of the G23S BglB mutation. Panel A shows that S23 and E26, which forms a hydrogen bond with the introduced serine, are located in a loop between a beta sheet and an alpha helix containing Q19. Panel B shows a zoomed view of the S23 residue site. Panel C shows that Q19 is positioned adjacent to the substrate, which is highlighted in brown. Residues of interest are highlighted in purple and blue, and hydrogen bonds in panels A and C are represented by a light blue dashed line.

It was hypothesized that relative to the wild-type (WT), the β-glucosidase G23S mutant would exhibit increased binding affinity (decreased K_m_), similar turnover number (k_cat_), increased catalytic efficiency (k_cat_/K_m_), increased maximum velocity (V_max_), and slightly reduced thermal stability (T_50_) based on the predicted additional hydrogen bond, prior D2D data on the G23A mutation, and a moderately increased Foldit score.

## RESULTS

To ensure successful transformation of the mutant DNA, Kunkel product transformation was compared to a ssDNA control plate (7). There were 860 colonies estimated on the Kunkel plate and 330 colonies on the control. The transformation efficiency was 2.6. To confirm proper introduction of the mutation into the plasmid, sequence analysis was performed. Colony #1 showed the best results, as shown in Figure 4, and was used for subsequent experiments. The quality scores of the bases in the desired mutant sequence were (61, 58, 58), indicating high base call accuracy. These data confirm the successful introduction of the mutation; proteins were then expressed.

**Figure 4.**
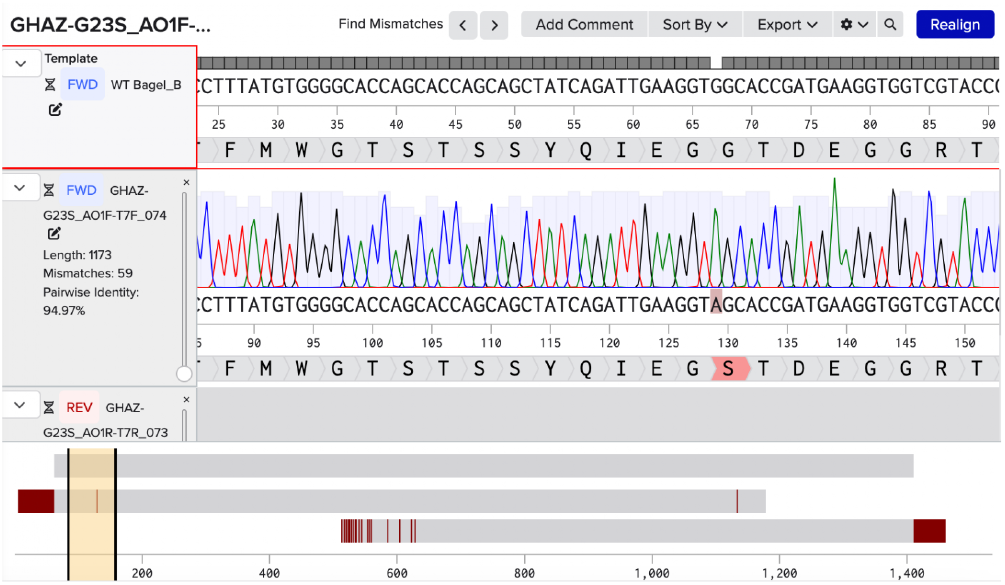
The DNA sequences of wild-type (WT) and G23S mutant BglB proteins were compared and the correct mutation in the colony was confirmed. To confirm the successful introduction of the mutation during the Kunkel reaction and subsequent *E. coli* (DH5*α*) transformation, the BglB region of the mutant plasmid was sequenced and compared to the WT sequence. For sequencing preparation, two PCR tubes (one with forward primer, one with reverse) were prepared for each colony. After the samples were submitted for sequencing, each sequence file (.ab1 format) contained the raw nucleotide string and associated chromatogram data. Four G23S mutation sequence files were compared to the WT sequence. The chromatogram data in the displayed file correctly substitutes S for G in the 23 position. The mutation is highlighted in red.

To determine proper protein purification, an SDS-PAGE gel was run for the mutant, WT, and flowthrough (FT) from column chromatography. The mutant showed up on the gel at ∼45 kDa as shown in Figure 5. While there was some contamination, it was not substantial. Also, some protein showed up in the FT, likely because all samples contained large amounts of protein. To determine protein concentration, an A280 assay was performed. As shown in Table 1, the mutant absorbance was 1.9148 and the WT absorbance was 1.2055 (a different WT sample with an A280 of 0.2405 was used in the kinetics assay). The mutant concentration was 0.8293 mg/mL and the WT concentration was 0.5615 mg/mL. These protein expression and purification results demonstrate successful purification; the protein was used for downstream assays.

**Table 1.** The protein concentration of WT and G23S BglB were determined to be sufficient for downstream assays. Concentration was ascertained from the A280 value, extinction coefficient, pathlength, and molecular weight. After protein purification, A280 values were determined using a Nanodrop spectrophotometer. The Nanodrop was set to the Protein A280 setting to read values at 280 nm. The Beer-Lambert Law (A = εbc) was then used to determine WT and G23S concentration in M. The pathlength value was 1 cm, and the extinction coefficients were computed using the Expasy ProtParam tool. The ProtParam tool was also used to calculate the molecular weights in Da (g/mol), which were used to express protein concentration in terms of mg/mL. The yield indicates sufficient protein present for downstream kinetic and thermal assays.

| BglB variant | A280 value (M*cm) | Protein concentration (mg/mL) |
| --- | --- | --- |
| WT | 1.2055 | 0.5615 |
| G23S | 1.9148 | 0.8293 |

**Figure 5.**
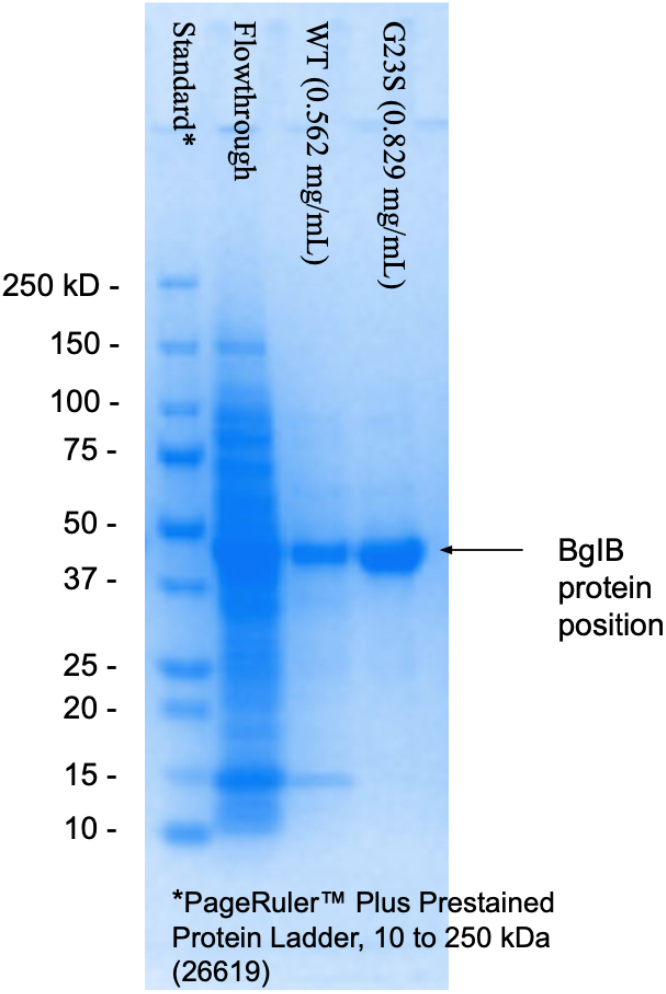
The wild-type (WT) and G23S mutant BglB proteins were successfully purified to homogeneity. *E. coli* BL21(DE3) cells expressing the WT and G23S mutant BglB were induced with IPTG, harvested, and lysed by sonication. Soluble protein fractions were isolated by centrifugation and eluted from a Ni-NTA resin column. GFP was also eluted to ensure the general protein elution process was successful. Samples from the flowthrough (FT), WT, and G23S fractions were separated on a 4-12% SDS-PAGE gel and visualized with Coomassie Brilliant Blue staining for analysis. Both WT and G23S lanes displayed a prominent band right below the 50 kDa band (∼45 kDa). While there was some contamination in the mutant and WT lanes, it was not substantial; while the FT lane included protein, that was likely because there was a large quantity of protein in the experiment overall, as shown in the WT and G23S lanes. The arrow represents the molecular weight of the BglB protein.

To determine kinetic parameters for WT and G23S BglB, a kinetic assay was run, Michaelis-Menten plots were generated by the D2D platform (Figure 6), and values were derived and shown in Table 2. Relative to the WT, the mutant K_m_ was approximately the same (7.94 ± 0.50 mM vs 7.36 ± 0.85 mM in WT), k_cat_ increased by almost 2-fold (1.28 × 10^3^ ± 24.2 min^-1^ vs 6.80 × 10^2^ ± 22.7 min^-1^ in WT), k_cat_/K_m_ also increased (161 min^-1^⋅mM^-1^ vs 92.4 min^-1^⋅mM^-1^ in WT), and V_max_ increased by almost 14-fold (1.52 × 10^-4^ M⋅min^-1^ vs 1.09 × 10^-5^ M⋅min^-1^ in WT).

**Table 2.** Kinetic values generated by the D2D platform show that G23S exhibits a similar K_m_ and increased k_cat_, k_cat_/K_m_, and V_max_ compared to the WT. Kinetic data was cleaned and filtered in Excel then uploaded to the D2D database to calculate K_m_, k_cat_, k_cat_/K_m_, and V_max_ values for the WT and G23S BglB enzymes. The trends generally align with the hypothesis: increased catalytic efficiency (k_cat_/K_m_) was observed. However, the hypothesis predicted increased binding affinity (decreased K_m_) would drive the enhanced catalytic efficiency; in the experiment, a higher k_cat_ improved the catalytic efficiency.

| Protein variant | $K_m$ (mM) | $k_{cat}/K_m$ (min<sup>-1</sup>·mM<sup>-1</sup>) | $k_{cat}$ (min<sup>-1</sup>) | $V_{max}$ (M/min) |
| --- | --- | --- | --- | --- |
| WT | $7.36 \pm 0.85$ | 92.4 | $6.80 \times 10^2 \pm 22.7$ | $1.09 \times 10^{-5}$ |
| G23S | $7.94 \pm 0.50$ | 161 | $1.28 \times 10^3 \pm 24.2$ | $1.52 \times 10^{-4}$ |

**Figure 6.**
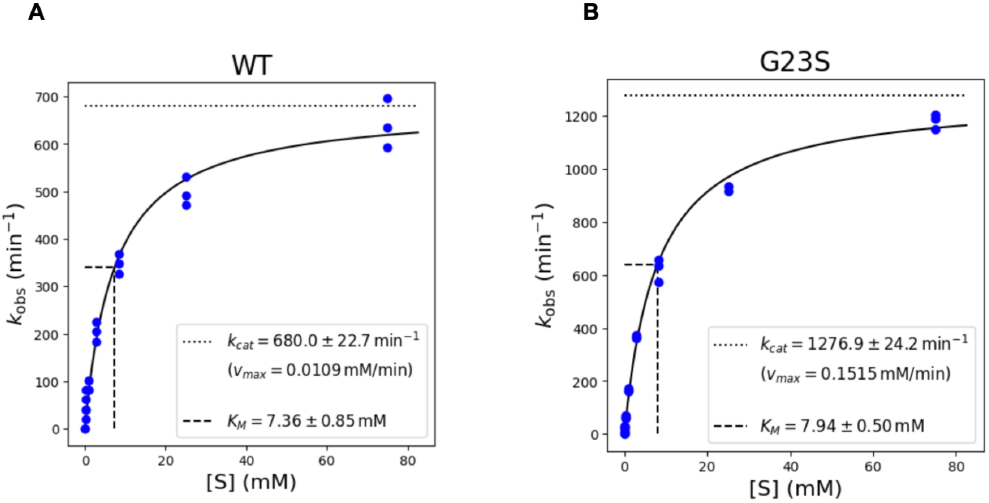
Michaelis-Menten kinetics curves of WT (panel A) and G23S (panel B) BglB demonstrated both enzyme variants followed typical saturation trends. Based on information from A280 calculations and a color change assay using the 4-nitrophenyl-β-D-glucopyranoside (pNP-Glc) substrate stock, WT and G23S protein were both diluted 1:100. Substrate concentrations varied from 0 mM to 75 mM according to a 3X serial dilution. A kinetic assay was performed using an MPM6 plate reader that measured absorbance at 420 nm every minute for 15 minutes. The color change of pNP-Glc was monitored in 96-well plates. Data was cleaned and filtered in Excel, then uploaded to the D2D database. The curves demonstrate that both BglB variants follow typical saturation trends; G23S has a higher V_max_ and similar K_m_ compared to the WT.

To determine enzyme stabilities, thermal assays were run, data was uploaded to the D2D database, and stability plots were generated (Figure 7). The T_50_ values of WT and G23S BglB, respectively, were 41.3 °C and 39.9 °C.

**Figure 7.**
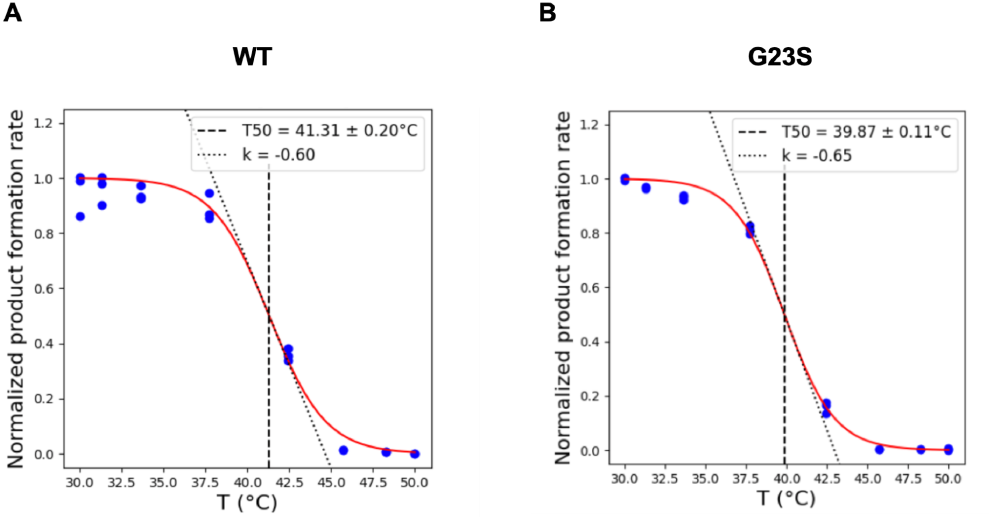
Thermal stability plots of WT (panel A) and G23S (panel B) BglB demonstrated that the T_50_ value of G23S BglB is slightly below that of the WT. As was done for the kinetics assay, based on information from A280 calculations and a color change assay using the 4-nitrophenyl-β-D-glucopyranoside (pNP-Glc) substrate stock, WT and G23S protein dilutions were determined. WT was diluted 1:25 and G23S was diluted 1:50. Protein and pNP-Glc substrate stocks were added to separate 96-well plates. Using a thermal cycler, aliquots of enzyme were exposed to a temperature gradient of 30–50 °C. An assay was performed using an MPM6 plate reader that measured absorbance at 420 nm every minute for 15 minutes. The color change of pNP-Glc was monitored in 96-well plates. Data was cleaned and filtered in Excel, then uploaded to the D2D database, which generated plots. The T_50_ of WT was 41.3 °C and that of G23S was 39.9 °C. Thus, the T_50_ difference was -1.4 °C, demonstrating that the WT was slightly more stable than the mutant.

## DISCUSSION

Based on Foldit modeling (Figure 2), the G23S mutation introduces a hydrogen bond while largely preserving overall enzyme conformation. The slight increase in Foldit score suggests modest global destabilization, while the structural model indicates potential improvements in local stability and substrate alignment. We therefore hypothesized that G23S would exhibit increased binding affinity (decreased K_m_), catalytic efficiency (k_cat_/K_m_), and maximum velocity (V_max_). Experimentally, binding affinity remained unchanged (similar K_m_), while an increase in turnover number (k_cat_) led to an increase in catalytic efficiency (k_cat_/K_m_) and maximum velocity (V_max_). Differences in V_max_ may partially reflect differences in enzyme concentration, so k_cat_ is the true comparison point. The increase in catalytic efficiency was driven by improved turnover rather than enhanced substrate binding. Based on the higher mutant Foldit score and prior D2D data on G23A, a slight decrease in thermal stability (T_50_) was predicted. (3) Experimentally, G23S exhibited a moderately reduced T_50_ (39.9 °C vs 41.3 °C in WT). Overall, the results are broadly consistent with the hypothesis.

Structurally, the serine in G23S forms a hydrogen bond with E26 within a loop downstream of an α-helix that contains Q19 (Figure 3). This interaction, along with the reduced conformational flexibility from mutating glycine, may alter local backbone geometry and residue interactions. As shown in Figure 3C, Q19 is located proximal to the substrate, indicating that the mutation may indirectly influence substrate orientation.

Local structural reorganization may promote a more favorable configuration for catalysis, potentially by enhancing transition state stabilization (9), although further experimental analysis is needed to validate this proposed mechanism.

The increased Foldit score is consistent with the observed decrease in thermal stability, which may reflect increased steric crowding introduced by the larger side chain relative to glycine. To further investigate steric constraints, future studies could utilize Partial Least Squares (PLS) models of ATR-IR and Raman Spectra to compare the secondary structures of the WT and G23S (10).

At the time of this publication, the only other D2D-characterized mutation at residue 23, G23A, exhibits decreased K_m_, similar k_cat_, increase in k_cat_/K_m_, and slightly reduced T_50_ relative to the WT. Although G23A does not introduce hydrogen bonding capability, it likely reduces conformational flexibility associated with glycine. In contrast, G23S introduces both steric and hydrogen bonding changes. G23S lacks binding affinity changes and uniquely increases k_cat_, suggesting a distinct mechanism. These results indicate that mutations at residue 23 can enhance catalytic efficiency while slightly decreasing stability.

Further studies could examine G23D, which introduces both steric bulk and additional hydrogen bonding capability compared to G23S. While this mutation may further contribute to steric crowding and decreased stability, its increased polarity could improve catalysis unless steric or electrostatic effects disrupt substrate binding.

Methodological aspects may also influence experimental results and interpretation of kinetic and thermal values. Synthetic substrates can yield kinetic parameters that are inconsistent with those obtained from natural substrates (11); thus, future studies should include comparisons with natural oligosaccharide substrates. In addition, incorporation of biological replicates would improve the reliability of measured kinetic and thermal parameters.

## CONCLUSION

The G23S mutation resulted in a similar K_m_, an approximately 2-fold increase in k_cat_ and k_cat_/K_m_, an almost 14-fold increase in V_max_, and a slight decrease in T_50_. Overall, experimental findings support the hypothesis, with the exception that the increase in catalytic efficiency was driven by a higher turnover number rather than an increased substrate binding affinity. Our model suggests a link between these kinetic changes and the hydrogen bond between S23 and E26, which may alter residue interactions and substrate orientation. The observed decrease in thermal stability is consistent with the steric constraints introduced by the larger side chain of serine relative to glycine. These findings suggest that substitutions at residue 23 can notably increase catalytic efficiency while modestly reducing stability, as well as highlight the sensitivity of enzyme function to point mutations distal from the active site.

## METHODS

All experimental procedures were performed according to standard protocols adapted from the Design2Data with β-glucosidase Lab Manual & Workflow (Siegel Laboratory, UC Davis, 2023) (8).

### Foldit Modeling and Analysis

The β-glucosidase (BglB) enzyme from Paenibacillus polymyxa was analyzed in silico using Foldit, a graphic user interface (GUI) and online gaming software that provides intermolecular modeling analysis features. (1) The G23S mutant version was modeled in Foldit and was selected primarily due to increased hydrogen bonding capabilities through the -OH group on serine. In addition, the Design to Data (D2D) database had data on one similar mutation, G23A, predicting a large increase in kcat/K_m_ due to a decrease in K_m_ with the same thermal stability.

### Mutagenesis

The Kunkel Mutagenesis technique was used to introduce our site-specific mutation into a plasmid. A 33 base pair oligonucleotide, identical to the target region except for at the mutation site, was synthesized. The full oligonucleotide sequence, 5’ to 3’, was ACCACCTTCATCGGTGCTACCTTCAATCTGATA. The codon responsible for mutating the target amino acid was GCT. The mutant oligonucleotide was phosphorylated by T4 Polynucleotide Kinase (T4PNK) at the 5’ end, annealed to single-stranded plasmid DNA, polymerized by T7 Polymerase (Cat# M0274L) to introduce the mutation into the plasmid, and ligated by T4 Ligase (Cat# M0202L) to make the plasmid circular. The reactions were completed as per manufacturer’s protocol. The Kunkel Product was incubated for 1 hour at room temperature (22.0 °C) then stored at -20 °C.

### DH5α Transformations

Then, chemically-competent *Escherichia coli* DH5αcells (NEB C2987H) were transformed using standard heat-shock transformation protocol, with the exception that the RPM of the shaking incubation during cell recovery was 190 (cells were incubated for 1 hour at 37 °C and 190 RPM). Following the protocol ranges, 2 µL of Kunkel product and 20 µL of DH5α cells were pipetted. Culture was pipetted onto LB agar plates with 50 µg/mL Kanamycin. Once dry, plates were incubated overnight at 37 °C. Labeled culture tubes were also prepared, with the exception that we prepared four instead of three tubes because our mutation occurred in the first 30 nucleotides of the sequence. By increasing the number of tubes, we increased our probability of a successful transformation that isolated a colony with the mutation, as well as high quality, accurate sequencing data.

### Overnight Cultures

Then, the transformed bacteria were grown to generate a sufficient quantity of plasmid DNA for downstream applications. One colony per tube was placed in four 15-mL round-bottom culture tubes with LB and Kanamycin growth media. The tubes were placed in a shaking incubator and incubated overnight at 37 °C.

### Plasmid Extraction

Transformation efficiency was analyzed using a ssDNA blank as a reference. The mutant plasmid was isolated from a 15 mL overnight culture using the NEB Monarch Plasmid Miniprep Kit according to the manufacturer’s instructions, with the exception that four instead of three samples were processed in parallel, as four tubes were prepared.

### DNA Concentration Assay (A260)

To confirm the success of the Miniprep plasmid isolation and assess DNA purity, an A260 assay was performed on a Nanodrop spectrophotometer. For each of the four samples, two measurements were taken, and the average value was used for plasmid sequence preparation concentrations.

### Plasmid Sequencing Preparation

Sequencing was performed using Sanger Sequencing (Applied Biosystems, 3730xL DNA Analyzers). Isolated plasmid DNA, 10 µM ‘T7 forward’ or ‘T7 reverse’ primers, and nuclease-free H_2_O, were prepared and placed into 0.5 mL conical base screw cap microcentrifuge tubes (purple lid), with the exception that no nuclease-free H_2_O was added for calculated DNA concentrations over 17.2 µL. The conical base tubes were used as opposed to the sequencing PCR tubes because we submitted four samples instead of the standard three. Thus, a total of eight tubes were submitted for sequencing (four with ‘T7 forward’ and four with ‘T7 reverse’ primers).

### Sequencing Results: Data Analysis

The sequence chromatogram data was stored on Benchling. Forward and reverse sequence reads were aligned with and compared to the WT BglB sequence. The plasmid with the read that best covered the G23S mutation out of the four total was used for downstream experiments.

### Chemically Competent BLR Transformation

1 µL of Mutant BglB, WT BglB, and GFP plasmids were then each transformed into 20 µL of BL21(DE3) chemically competent *E. coli* (Thermo Scientific™ EC0114) using standard heat-shock transformation protocol. A culture tube was also prepared, and the culture was plated on a Kanamycin agar plate for overnight growth at 37 °C.

### Protein Production: Growth and Expression of Overnight Cultures

Transformed bacteria were cultured to increase cell numbers and ensure enough protein for downstream applications. Breathe-Easy^®^ sealing membrane (Cat No. Z380059) was used to seal the cultures, which were incubated at 37 °C with shaking for 24 hours. Then, expression cultures were prepared using the engineered pET expression system. The 50 mL culture was induced with a final concentration of 1 mM of isopropyl β-D-1-thiogalactopyranoside (IPTG). IPTG-containing cultures were incubated at 18 °C with shaking overnight. The pellets were stored at 20 °C for downstream applications.

### Protein Purification

Mutant enzymes, along with the WT protein and GFP, were purified using immobilized metal affinity chromatography (IMAC) to isolate the BglB enzyme from contaminants. Cell lysis and clarification was performed using the freeze-thaw cycles and the BugBuster detergent mix. Then, Ni-NTA columns were used for isolation. The His-tagged BglB enzyme was purified using the column, and EDTA solution was used for elution according to standard affinity chromatography protocol, with the exception that samples were recycled with eluate three times because the GFP (visual representation of elution efficacy) did not elute from the column after two rounds of eluate (150 mL) and resuspension.

### Quantification of Protein Yield (A280)

The Nanodrop spectrophotometer was used to estimate protein concentration by A280. Glycerol was added to the protein for stabilization.

### SDS-PAGE

A Sodium Dodecyl Sulfate-Polyacrylamide Gel Electrophoresis (SDS-PAGE) was used to further verify protein purity. 4x loading dye (Lamelli sample buffer without added β-mercaptoethanol), protein gel (4–20% Mini-PROTEAN® TGX Stain-Free™ Protein Gels, 15 well, 15 µl #4568096), Precision Plus Protein™ All Blue Prestained Protein Standards (Catalog #1610373) marker, 1X MES running buffer, and Coomassie Blue protein gel stain were used. Mutant protein, WT protein, flowthrough, and PageRuler™ Plus Prestained Protein Ladder were loaded onto the gel. The gel was run and imaged. Mutant and WT protein concentrations were calculated using the Beer-Lambert Law, previously determined A280 values, a pathlength of 1cm, and molar extinction coefficients from the Expasy ProtParam tool.

### Kinetic Assay

A kinetic assay was performed to compare mutant catalytic efficiency to that of the WT. The color change of 4-nitrophenyl-β-D-glucopyranoside (pNP-Glc), a synthetic substrate, was monitored in 96-well plates. The substrate plate was a half-area 96-well plate, and the protein plate used was Corning 96-Well Clear-Bottom Half-Area Microplate, #3884. The Michaelis-Menten kinetic model was used to quantify the rate of color change. Substrate concentrations in a range of 0 mM to 75 mM (75 mM, 25 mM, 8.33 mM, 2.78 mM, 0.93 mM, 0.31 mM, 0.10 mM, 0.00 mM) were varied by a 3X serial dilution. Purified proteins were diluted (1:100 for both G23S and WT) based on A280-derived calculations, with the exception that a different WT (yield = 0.11 mg/mL was used). 25 µL of diluted purified protein was added to each well across three adjacent columns. To begin the assay, 75 µL of solutions from the substrate plate were transferred to the protein plate such that they were overlaid (i.e., A1 in the substrate plate to A1 in the protein plate). In total, the proteins were diluted 1:100, 1:1.36 (addition of glycerol), and 1:4 (addition of substrate). The final concentration of WT was 2.0 × 10_-4_ mg/mL and G23S was 1.5 × 10_-3_ mg/mL. An MPM6 plate reader was used to measure absorbance at 420 nm every minute for 15 minutes.

### Thermal Stability Heat Challenge Assay

A thermal stability heat challenge assay was performed to determine the T_50_ value. Purified mutant protein was diluted (1:50) and WT protein was diluted (1:25). 50 µL of protein was added to each well of a half-area 96-well plate in triplicates. 75 µL of 10 mM substrate solution was added to a Corning 96-Well Clear-Bottom Half-Area Microplate, #3884. In total, the proteins were diluted 1:50 or 1:25, 1:1.36 (addition of glycerol), and 1:4 (addition of substrate). The final concentration of WT was 4.1 × 10^-3^ mg/mL and G23S was 3.1 × 10^-3^ mg/mL. Using a thermal cycler, aliquots of the enzyme were exposed to a variety of temperatures to create a temperature gradient of 30–50 °C (50.0 °C, 48.3 °C, 45.7 °C, 42.4 °C, 37.7 °C, 33.6 °C, 31.3 °C, 30.0 °C). Activity was determined using the same pNP-Glc and 96-well substrate and protein plate method as the kinetics assay, with the exception that 25 µL of protein was transferred instead of 75 µL of substrate solution, according to standard protocol.

### Data Analysis

Last, data was cleaned, filtered, and submitted to the D2D Database. Results were evaluated.

## ACKNOWLEDGMENTS

This work was supported by the Siegel Laboratory at University of California, Davis, and the Department of Molecular Biochemistry & Biophysics at Yale University.

